# Purge Haplotigs: Synteny Reduction for Third-gen Diploid Genome Assemblies

**DOI:** 10.1101/286252

**Authors:** Michael J Roach, Simon Schmidt, Anthony R Borneman

## Abstract

Recent developments in third-gen long read sequencing and diploid-aware assemblers have resulted in the rapid release of numerous reference-quality assemblies for diploid genomes. However, assembling highly heterozygous genomes is still facing a major problem where the two haplotypes for a region are highly polymorphic and the synteny is not recognised during assembly. This causes issues with downstream analysis, for example variant discovery using the haploid assembly, or haplotype reconstruction using the diploid assembly. A new pipeline—Purge Haplotigs—was developed specifically for third-gen assemblies to identify and reassign the duplicate contigs. The pipeline takes a draft haplotype-fused assembly or a diploid assembly, and read alignments to produce an improved assembly. The pipeline was tested on a simulated dataset and on four recent diploid (phased) *de novo* assemblies from third-generation long-read sequencing. All assemblies after processing with Purge Haplotigs were less duplicated with minimal impact on genome completeness. The software is available at https://bitbucket.org/mroachawri/purge_haplotigs under a permissive MIT licence.

## Background

Recent advances in third-generation single-molecule sequencing have enabled *de novo* genome assemblies that have extremely high levels of contiguity and completeness (Badouin et al., 2017, Jarvis et al., 2017, Loman et al., 2015). Furthermore, recent advances in ‘diploid aware’ genome assemblers have considerably improved the quality of highly heterozygous diploid genome assemblies (Chin et al., 2016, Korlach et al., 2017). Diploid-aware assemblers such as FALCON and Canu are available that will produce a haplotype-fused representation of a diploid genome (Chin et al., 2016, Koren et al., 2017), and some assemblers such as FALCON Unzip and Supernova will go further to produce large phase blocks where both parent alleles are represented separately (Chin et al., 2016, Weisenfeld et al., 2017). For FALCON Unzip assemblies, which are the focus of this study, phasing occurs on the assembly graph to produce ‘primary contigs’ (the haploid assembly) and associated ‘haplotigs’, which together with the primary contigs form the diploid assembly.

Regions of very high heterozygosity still present a problem for *de novo* genome assembly (Kajitani et al., 2014, Safonova et al., 2015, Vinson et al., 2005). In this situation, once a pair of allelic sequences exceeds a certain threshold of nucleotide diversity, most algorithms will assemble these regions as separate contigs, rather than the expected single haplotype-fused contig (Pryszcz et al., 2014, Small et al., 2007). The presence of these syntenic contigs in a haploid assembly is problematic for downstream analysis (Olson et al., 2015). In the case of producing a diploid assembly, while both alleles may be present, steps are still required to identify the syntenic contig pairings.

Several tools have attempted to deal with this problem. The HaploMerger2 toolkit (Huang et al., 2017) and Redundans assembly pipeline (Pryszcz and Gabaldon, 2016) were designed to produce haplotype-fused assemblies from short-read sequences. However, both include steps that would not generally be employed for finishing an already highly contiguous long-read based assembly. Furthermore, resolving the haplotype sequences and producing a phased assembly has proven to be advantageous (Schwessinger et al., 2018, VanBuren et al., 2018). Scripts available for use with long-read assemblies include; get_homologs.py, which uses sequence alignments to identify homologues (Concepcion, 2016) and HomolContigsByAnnotation, which uses gene annotations to match syntenic regions (Kingan, 2016). Each has its unique strengths and drawbacks, but both suffer from requiring manual reassignment of contigs by the user.

The aim of this study was to develop a new pipeline that could quickly and automatically identify and reassign syntenic contigs specifically in assemblies produced with single-molecule long-read sequencing technology. Purge Haplotigs is designed to be easy to install and requires only three commands to complete. It will work on either the haploid assembly to produce a de-duplicated haploid assembly, or on the diploid assembly to produce a de-duplicated haploid assembly and an improved diploid assembly.

### Implementation

The Purge Haplotigs pipeline is outlined in Figure 1. The pipeline requires two input files: a draft assembly in FASTA format, and an alignment file of reads mapped to the assembly in BAM format. The input draft assembly can be either the haploid or diploid assembly. For the aligned reads, the pipeline works best when the long-reads that were used for generating the assembly are mapped, but it will also work using short reads. A ‘random best’ alignment should be used for multi-mapping reads and the library should be one that produces an unbiased flat read-coverage.

**Figure 1:**
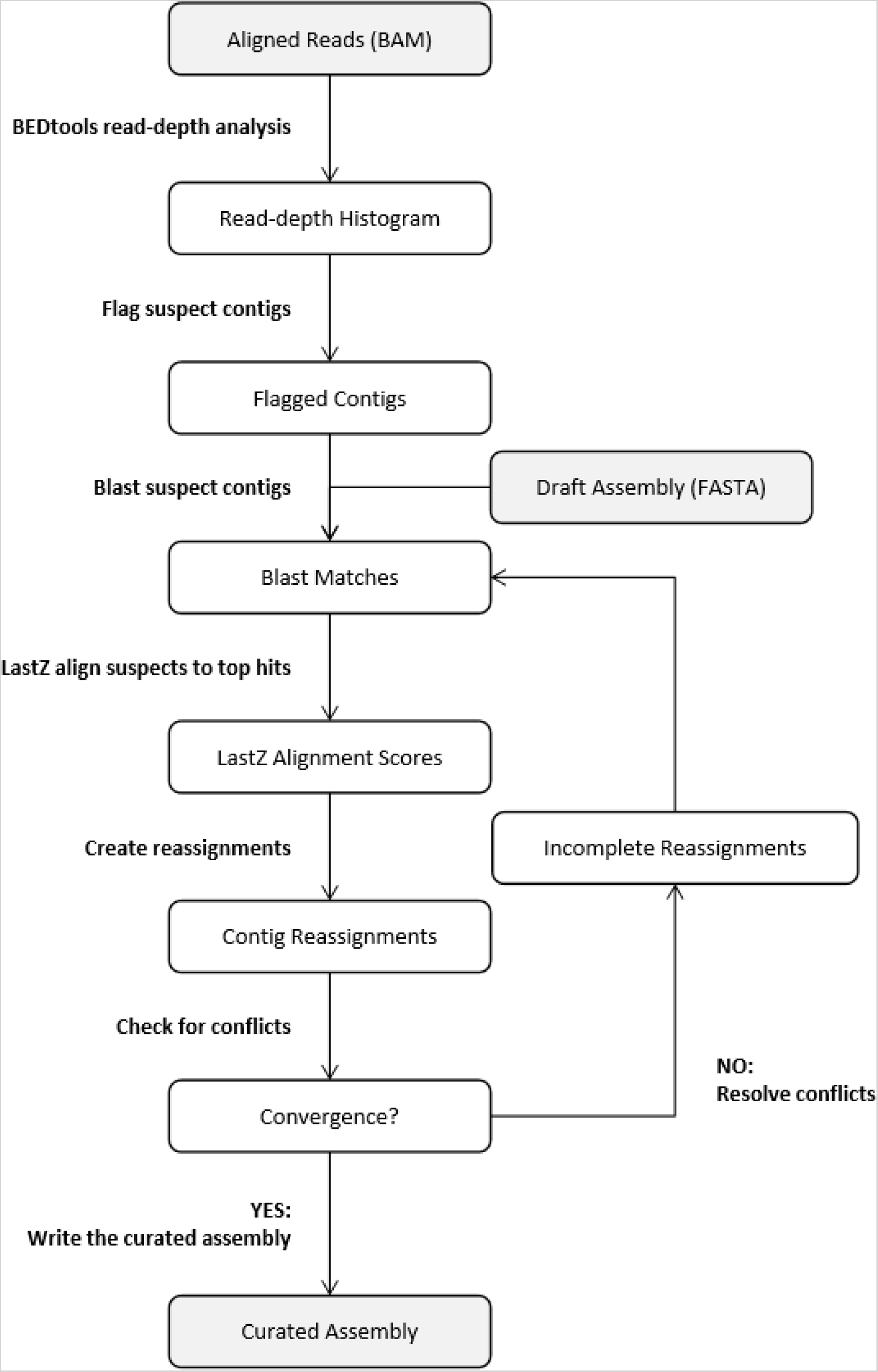
Flow chart for the Purge Haplotigs pipeline.

### Read-depth analysis

The first stage involves a read-depth analysis of the BAM file using BEDtools (Quinlan and Hall, 2010). A read-depth histogram is initially produced for the assembly. For collapsed haplotype contigs the reads from both alleles will map, whereas if the alleles have assembled as separate contigs the reads will be split over the two contigs, resulting in half the read-depth. We exploit this to flag contigs that are likely to be haplotigs.

For a haploid assembly, a bimodal distribution should be observed if duplication has occurred (Figure 2). The left peak results from the duplicated regions and the right peak at twice the read-depth results from regions that are properly haplotype-fused. For a diploid assembly, as the entire assembly should be duplicated, the second peak should only be very small or not visible at all. The user chooses three cut-offs to capture the two peaks and the pipeline then calculates a breakdown of the read-depth proportions for each contig. Contigs with a high proportion of bases within the ‘duplicated’ range for read-depth are flagged for further analysis. For a diploid assembly, as both haplotypes should be present, most of the contigs would be expected to be flagged for further analysis.

**Figure 2:**
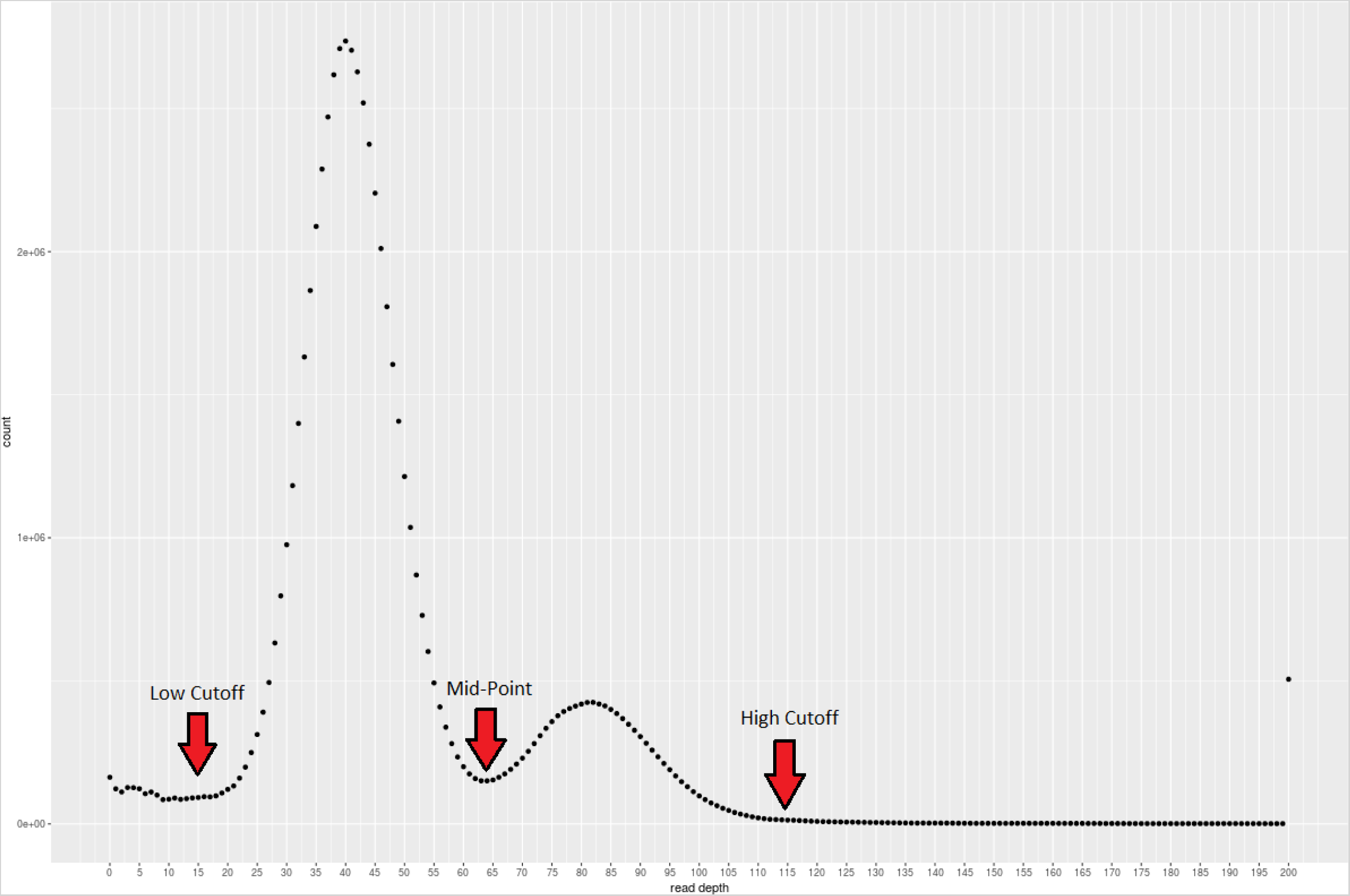
Example read-depth histogram produced by Purge Haplotigs. This example for *C. pyxidata* was produced using PacBio RS II reads mapped to the diploid assembly. Example cut-offs are indicated for use with the second stage of the pipeline.

Contigs that have a majority of their bases displaying a read-depth outside of the defined bounds (abnormally low or high coverage) are further flagged for removal with the assumption that they are artefactual. It should be noted that contigs from organelle DNA sources may have a much higher read-depth than the rest of the genome, as such these may appear with the artefactual contigs after processing with Purge Haplotigs.

### Identification and assignment of homologous sequences

Contigs that were flagged for further analysis according to read-depth are then subject to sequence alignment to attempt to identify synteny with its allelic companion contig. All flagged contigs therefore undergo a BLAST search (Camacho et al., 2009) against the entire assembly to identify discrete regions of nucleotide similarity. Chained alignments are then calculated using LASTZ (Harris, 2007) for each flagged contig against its BLAST best hit(s). Using these data Purge Haplotigs then calculates both the total portion of the flagged contig that aligns at least once (alignment score) and the sum of all alignments (max match score) between the flagged contig and its best hit contigs. The alignment score is used to determine if each flagged contig should be reassigned as a haplotig, while the max match score determines if it should instead be labelled as repetitive. The max match score is intended to highlight problematic contigs such as collapsed repeats. It should be noted that highly repetitive genomic regions, such as centromeres and telomeres, may also be labelled as repetitive contigs. Conflicts may arise where haplotigs are nested, overlap, or are comprised of mostly repetitive sequence. This can cause individual contigs to be both flagged for reassignment and flagged as a reference for reassigning another contig. Where this occurs, the pipeline will only purge the contig that is most likely to be a nested haplotig or collapsed repeat. Because of this the LASTZ alignments, scoring, and conflict resolution occurs iteratively until no more conflicts occur and no more contigs meet the conditions for reassignment as a haplotig.

### Outputs

Purge Haplotigs produces three FASTA format files for the curated assembly: the curated contigs, the contigs reassigned as haplotigs, and the contigs reassigned as artefacts. If the original input were a draft haploid assembly, then the curated contigs would represent the haploid assembly. Alternatively, if the original input were a draft diploid assembly then the curated contigs represent the haploid assembly, while the revised diploid assembly would consist of the combination of both the curated primary contigs and the reassigned haplotigs.

In addition to the FASTA output, Purge Haplotigs also produces several metrics to aid in the manual assessment of the automatic contig assignment function, including the production of dotplots juxtaposed with read-depth tracks for each reassigned and ambiguous contig. A data table is also produced which lists each contig reassignment and includes both the alignment and max match scores. Finally, a text file is produced to show the contig purging order for the situations in which conflicts were detected. This last file is particularly useful for producing dotplots for visualizing haplotig nesting and overlaps, as well as assessing any potential over-purging (for instance if the threshold for reassignment were set too low).

### Limitations

It should be noted that haplotype switching often occurs in the FALCON Unzip primary contigs between neighbouring phase blocks. The breaks in phasing usually occur for a reason and longer-range connectivity information is generally needed to completely reconstruct the two haplomes. As such Purge Haplotigs cannot resolve haplotype switching. Instead, it will only attempt to identify contigs that are syntenic and produce a de-duplicated representation of the genome.

## Results and Discussion

### Case Study

The Purge Haplotigs pipeline was first validated using a synthetic dataset (Additional File 1). However, to fully investigate the practical aspects and impact of synteny reduction, Purge Haplotigs was also tested on four draft FALCON Unzip assemblies. Assemblies for *Arabidopsis thaliana* (Cvi-0 × Col-0), *Clavicorona pyxidata* (a coral fungus), and *Vitis vinifera L. Cv*. Cabernet Sauvignon (grapevine) were sourced from Chin et al. (2016), and a fourth assembly for *Taeniopygia guttata* (Zebra finch) genome was sourced from Korlach et al. (2017). For each assembly, alignment files which consisted of PacBio RS II SMRT subreads mapped to each of the draft diploid assemblies, were generously provided by Pacific Biosciences.

### Methods

Assembly metrics were calculated using Quast v4.5 (Gurevich et al., 2013). Genome completeness, duplication, and fragmentation were predicted using BUSCO v3.0.1 (Simão et al., 2015). The MUMmer package v4.0.0 (Kurtz et al., 2004) was used to produce genome alignments and dotplots. Haploid assemblies were assessed for their performance using short read data. Suitable Illumina paired-end (PE) short reads were publicly available from the Short Read Archive (SRA) for *A. thaliana* Col-0 × Cvi-0 (SRA accessions: SRR3703081, SRR3703082, SRR3703105), *C. pyxidata* (SRA accession: SRR1800147), and *T. guttata* (SRA accession: ERR1013157). PE reads were downloaded and mapped using BWA-MEM v0.7.12 (Li, 2013) to the draft and curated haploid assemblies. Heterozygous SNPs were called using VarScan v2.3.9 (Koboldt et al., 2012), and read-coverage and SNP density were analysed using BEDtools v2.25.0 (Quinlan and Hall, 2010). The SNP density and read-depth histograms were visualized as Circos plots (Krzywinski et al., 2009). Detailed workflows for processing with Purge Haplotigs and subsequent analysis are available in Additional File 1.

### Assembly statistics

The removal of artefactual contigs resulted in the assemblies processed by Purge Haplotigs having 13–27 % fewer contigs (*A. thaliana* Table 1, Additional File 2). More importantly, a common problem with haploid assemblies contaminated by syntenic contigs, is that the final assembly size is significantly larger than the actual haploid genome size. The reassigning of redundant contigs by Purge Haplotigs reduced the total haploid assembly sizes for all four assemblies by 3.0–12.5 %. The draft FALCON Unzip haploid assembly for *A. thaliana* was 140 Mb, much larger than the current TAIR10 reference genome of 119 Mb (Lamesch et al., 2012). The Purge Haplotigs haploid assembly was 127 Mb, placing it close the expected haploid size. Likewise, the draft Cabernet Sauvignon haploid assembly was 591 Mb, much larger than the expected size of approximately 500 Mb for *V. vinifera* (Jaillon et al., 2007). After processing with Purge Haplotigs the improved assembly was reduced to 517 Mb.

**Table 1:** Assembly statistics for draft FALCON Unzip and Purge Haplotigs-processed *A. thaliana* assemblies.

|  | Primary Contigs |  | Haplotigs |  |
| --- | --- | --- | --- | --- |
|  | Original | Curated | Original | Curated |
| **Contigs** | 172 | 107 | 248 | 201 |
| **Largest contig** | 13 319 401 | 13 319 401 | 11 648 134 | 11 648 134 |
| **Total length** | 140 024 976 | 126 787 811 | 104 934 860 | 116 306 003 |
| **GC (%)** | 36.67 | 36.68 | 36.12 | 36.15 |
| **N50** | 7 960 654 | 7 979 657 | 6 920 133 | 4 634 947 |

**Table 2:** BUSCO statistics for draft FALCON Unzip and Purge Haplotigs-processed *A. thaliana* assemblies.

| **Haploid Assembly** |  | **FALCON Unzip** |  | **Purge Haplotigs** |  |
| --- | --- | --- | --- | --- | --- |
| **(Primary contigs)** |  | **#** | **%** | **#** | **%** |
| Total BUSCO groups searched |  | 1440 | 100.0 | 1440 | 100.0 |
| Complete BUSCOs |  | 1413 | 98.1 | 1408 | 97.8 |
| Complete and single-copy BUSCOs |  | 1324 | 91.9 | 1376 | 95.6 |
| Complete and duplicated BUSCOs |  | 89 | 6.2 | 32 | 2.2 |
| Fragmented BUSCOs |  | 5 | 0.3 | 9 | 0.6 |
| Missing BUSCOs |  | 22 | 1.5 | 23 | 1.6 |
| **Diploid Assembly** |  | **FALCON Unzip** |  | **Purge Haplotigs** |  |
| **(Primary + Haplotigs)** |  | **#** | **%** | **#** | **%** |
| Total BUSCO groups searched |  | 1440 | 100.0 | 1440 | 100.0 |
| Complete BUSCOs |  | 1414 | 98.2 | 1414 | 98.2 |
| Complete and single-copy BUSCOs |  | 70 | 4.9 | 70 | 4.9 |
| Complete and duplicated BUSCOs |  | 1344 | 93.3 | 1344 | 93.3 |
| Fragmented BUSCOs |  | 4 | 0.3 | 4 | 0.3 |
| Missing BUSCOs |  | 22 | 1.5 | 22 | 1.5 |
| **Phase Blocks** |  | **FALCON Unzip** |  | **Purge Haplotigs** |  |
| **(Haplotigs only)** |  | **#** | **%** | **#** | **%** |
| Total BUSCO groups searched |  | 1440 | 100.0 | 1440 | 100.0 |
| Complete BUSCOs |  | 1342 | 93.2 | 1397 | 97.0 |
| Complete and single-copy BUSCOs |  | 1313 | 91.2 | 1371 | 95.2 |
| Complete and duplicated BUSCOs |  | 29 | 2.0 | 26 | 1.8 |
| Fragmented BUSCOs |  | 5 | 0.3 | 4 | 0.3 |
| Missing BUSCOs |  | 93 | 6.5 | 39 | 2.7 |

### Synteny reduction and genome completeness

For the diploid assemblies, there were only minor differences comparing the draft and processed assemblies with respect to the predicted genome completeness and duplication, as indicated in the BUSCO analysis (*A. thaliana* Table 1, Additional File 2). For the haploid assemblies, the predicted level of duplication in the draft *C. pyxidata* and *T. gutatta* assemblies was relatively low at 3.7 % and 4.8 % respectively. The predicted duplication for the draft *A. thaliana* and Cabernet Sauvignon assemblies were higher at 6.2 % and 12.4 % respectively. The processed haploid assemblies contained between 40–74 % fewer duplicated BUSCOs than the draft haploid assemblies. Predicted genome completeness was minimally impacted. The *C. pyxidata* processed assembly contained 0.3 % more missing BUSCOs, but surprisingly the other processed assemblies contained up to 3.2 % *fewer* missing BUSCOs. Furthermore, the processed haplotigs contained 2.1–4.6 % fewer missing BUSCOs, suggesting that the haplotigs are themselves more complete representations of their genomes after processing with Purge Haplotigs.

### Phasing coverage

Proper identification of syntenic contig pairs results in improved phasing coverage of diploid assemblies. To assess if Purge Haplotigs provided improvements to this metric, pairwise alignments were performed between the primary contigs and haplotigs for both the draft and processed assemblies, and the total coverage of primary contigs by haplotigs was calculated (*A. thaliana* Figure 3; Additional File 3). For the *C. pyxidata* and *T. gutatta* assemblies the phasing coverage increased by 6.2 % and 7.9 % respectively. The two plant assemblies—which had higher predicted duplication— showed larger increases in phasing coverage of 12.8 % and 15.8 % for *A. thaliana* and Cabernet Sauvignon respectively.

**Figure 3:**
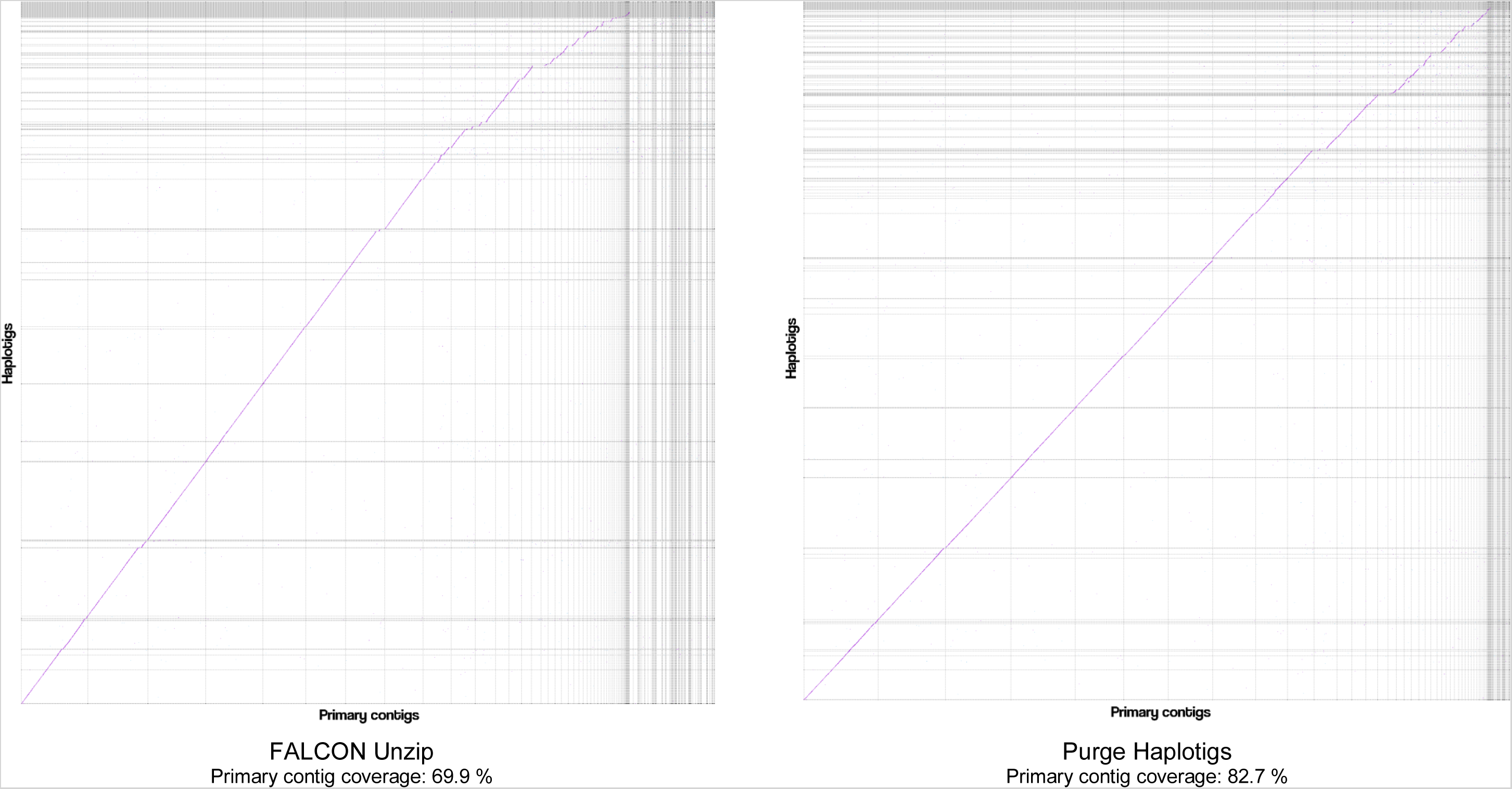
Dotplots for *Arabidopsis thaliana* assemblies. Haplotigs were aligned to primary contigs, filtered for one-to-one best alignments, coverage of the primary contigs by haplotigs calculated, and dotplots were laid out by longest alignments. Vertical gaps correspond to sequence in haplotigs that is not present in the primary contigs, and horizontal gaps correspond to sequence in the primary contigs not present in the haplotigs.

### Short-read performance

As mentioned previously, the erroneous presence of both syntenic contigs in a haploid assembly results in the presence of mapped regions displaying half the average read-depth and few (if any) heterozygous variant calls relative to the rest of the genome. To determine if the use short-reads for genomic analysis was improved after processing, combined read-depth and heterozygous SNP density plots were generated for both the draft and processed assemblies of *A. thaliana, C. pyxidata*, and *T. guttata* based upon the results from mapping illumina PE short read data (*A. thaliana* Figure 4; Additional File 4). The mapping rates of the processed assemblies only increased by 0.6–0.84 % compared to the draft assemblies. However, for *A. thaliana* there were approximately 14.5 % more heterozygous SNPs called for the processed assembly compared to the draft FALCON Unzip assembly. Likewise, there were 2.2 % and 12.5 % more heterozygous SNPs called for *T. gutatta* and *C. pyxidata* respectively.

**Figure 4:**
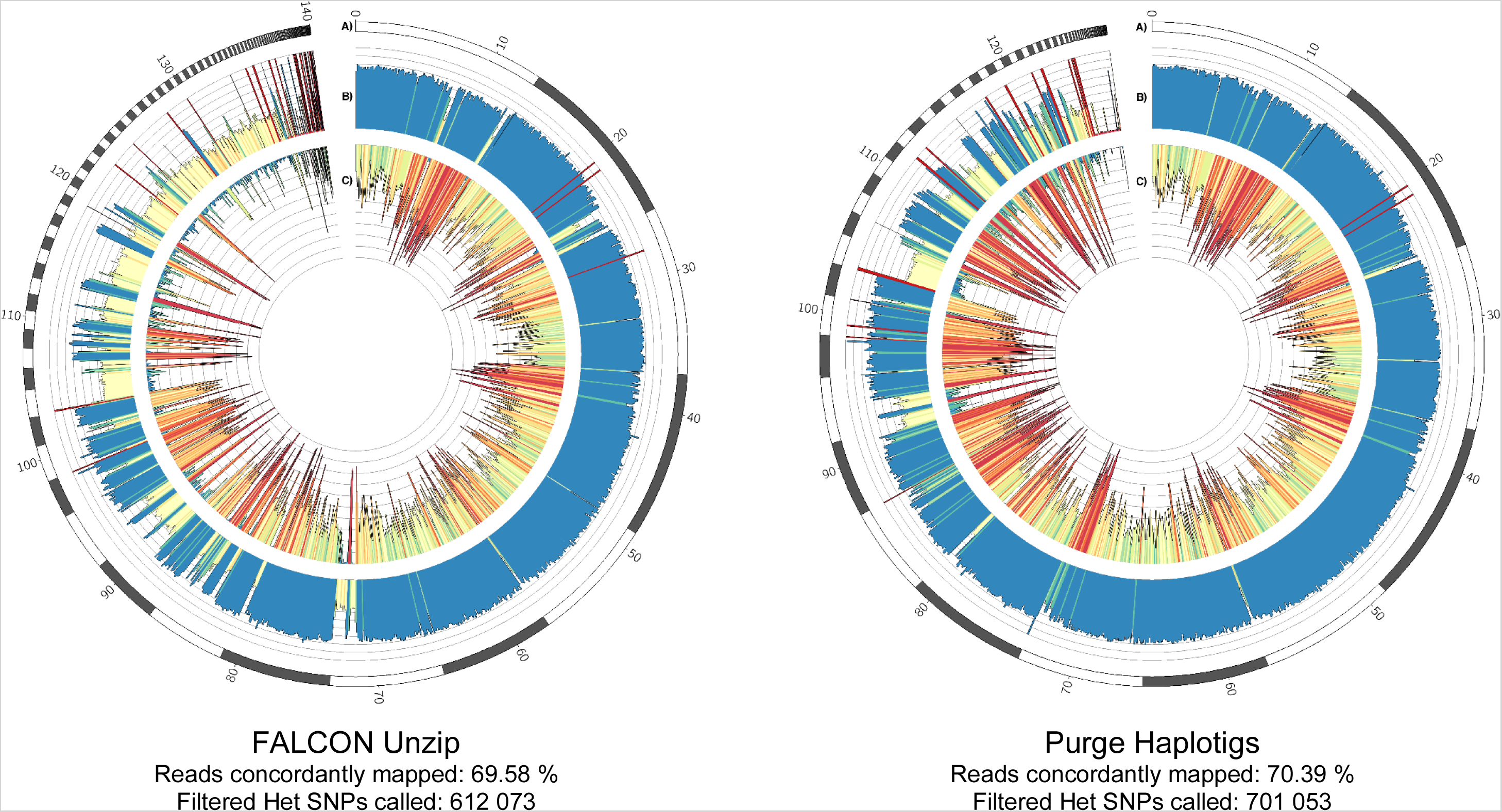
Circos plots for *Arabidopsis thaliana* haploid assemblies. Illumina PE reads were mapped and heterozygous SNPs were called for the draft FALCON Unzip assembly (LEFT) and the assembly curated with Purge Haplotigs (RIGHT). The tracks shown in the circos plots are: **A)** Contigs (ordered by length), **B)** Read-depth histogram (reads per genome window), and **C)** SNP density (SNPs per genome window).

## Conclusions

Purge Haplotigs is an effective tool for the early stages of curating highly heterozygous genome assemblies produced from third-generation long read sequencing. It can produce a mostly de-duplicated haploid representation of a genome which is important for downstream analysis such as variant discovery. Purge Haplotigs can also generate an improved diploid representation of a genome with more syntenic contigs identified and properly paired. This is particularly important for diploid assemblies, for instance if attempting to reconstruct parent haplomes.

## Availability and Requirements

**Project name:** Purge Haplotigs

**Project home page:** https://bitbucket.org/mroachawri/purge_haplotigs

**Operating system:** Linux (tested on Ubuntu 16.04 LTS)

**Programming language:** Perl

**Dependencies:** BEDTools, SAMTools, BLAST, LASTZ, Perl (with FindBin, Getopt::Long, Time::Piece, threads, Thread::Semaphore), Rscript (with ggplot2 and scales), GNU Parallel

**License:** MIT

**Restrictions:** None

## Abbreviations

PE: Paired End
SRA: Short Read Archive

## Acknowledgements

We would like to thank Sarah Kingan, Gregory Concepcion, Jason Chin and Pacific Biosciences for providing the BAM files for the assemblies and for helpful discussions.

## Supplementary Information

Additional File 1: Workflows for Purge Haplotigs and subsequent analysis.

- Workflows.pdf Additional File 2: Quast and BUSCO analysis results for all assemblies.
- Quast_BUSCO.xlsx Additional File 3: Circos Plots and mapping statistics for *C. pyxidata*, and *T. guttata*.
- Circos.pdf Additional File 4: Dotplots and coverage for *C. pyxidata, V. vinifera L. Cv*. Cabernet Sauvignon, and *T. guttata*.
- Dotplots.pdf

## Availability of Data

The simulated genome dataset is available at: https://doi.org/10.5281/zenodo.1042847. The dataset for the analysis described in this study of the draft and curated genome assemblies is available at: https://doi.org/10.5281/zenodo.1043619.

## References

Badouin, H., Gouzy, J., Grassa, C. J., Murat, F., Staton, S. E., Cottret, L., Lelandais-BrièRe, C., Owens, G. L., CarrèRe, S., Mayjonade, B., Legrand, L., Gill, N., Kane, N. C., Bowers, J. E., Hubner, S., Bellec, A., BéRard, A., BergèS, H., Blanchet, N., Boniface, M.-C., Brunel, D., Catrice, O., Chaidir, N., Claudel, C., Donnadieu, C., Faraut, T., Fievet, G., Helmstetter, N., King, M., Knapp, S. J., Lai, Z., Le Paslier, M.-C., Lippi, Y., Lorenzon, L., Mandel, J. R., Marage, G., Marchand, G., Marquand, E., Bret-Mestries, E., Morien, E., Nambeesan, S., Nguyen, T., Pegot-Espagnet, P., Pouilly, N., Raftis, F., Sallet, E., Schiex, T., Thomas, J., Vandecasteele, C., VarèS, D., Vear, F., Vautrin, S., Crespi, M., Mangin, B., Burke, J. M., Salse, J., MuñOs, S., Vincourt, P., Rieseberg, L. H. & Langlade, N. B. 2017. The sunflower genome provides insights into oil metabolism, flowering and Asterid evolution. Nature, 546, 148–152.

Camacho, C., Coulouris, G., Avagyan, V., Ma, N., Papadopoulos, J., Bealer, K. & Madden, T. L. 2009. BLAST+: architecture and applications. BMC Bioinformatics, 10, 421.

Chin, C. S., Peluso, P., Sedlazeck, F. J., Nattestad, M., Concepcion, G. T., Clum, A., Dunn, C., O’Malley, R., Figueroa-Balderas, R., Morales-Cruz, A., Cramer, G. R., Delledonne, M., Luo, C., Ecker, J.R., Cantu, D., Rank, D. R. & Schatz, M. C. 2016. Phased diploid genome assembly with single-molecule real-time sequencing. Nat Methods, 13, 1050–1054.

Concepcion, G. 2016. get_homologs.py [Online]. Available: https://github.com/PacificBiosciences/apps-scripts [Accessed 2017].

Gurevich, A., Saveliev, V., Vyahhi, N. & Tesler, G. 2013. QUAST: quality assessment tool for genome assemblies. Bioinformatics, 29, 1072–1075.

Harris, R. S. 2007. Improved pairwise alignment of genomic DNA, The Pennsylvania State University.

Huang, S., Kang, M. & Xu, A. 2017. HaploMerger2: rebuilding both haploid sub-assemblies from high-heterozygosity diploid genome assembly. Bioinformatics, 33, 2577–2579.

Jaillon, O., Aury, J. M., Noel, B., Policriti, A., Clepet, C., Casagrande, A., Choisne, N., Aubourg, S., Vitulo, N., Jubin, C., Vezzi, A., Legeai, F., Hugueney, P., Dasilva, C., Horner, D., Mica, E., Jublot, D., Poulain, J., Bruyere, C., Billault, A., Segurens, B., Gouyvenoux, M., Ugarte, E., Cattonaro, F., Anthouard, V., Vico, V., Del Fabbro, C., Alaux, M., Di Gaspero, G., Dumas, V., Felice, N., Paillard, S., Juman, I., Moroldo, M., Scalabrin, S., Canaguier, A., Le Clainche, I., Malacrida, G., Durand, E., Pesole, G., Laucou, V., Chatelet, P., Merdinoglu, D., Delledonne, M., Pezzotti, M., Lecharny, A., Scarpelli, C., Artiguenave, F., Pe, M. E., Valle, G., Morgante, M., Caboche, M., Adam-Blondon, A. F., Weissenbach, J., Quetier, F. & Wincker, P. 2007. The grapevine genome sequence suggests ancestral hexaploidization in major angiosperm phyla. Nature, 449, 463–7.

Jarvis, D. E., Ho, Y. S., Lightfoot, D. J., Schmöckel, S. M., Li, B., Borm, T. J. A., Ohyanagi, H., Mineta, K., Michell, C. T., Saber, N., Kharbatia, N. M., Rupper, R. R., Sharp, A. R., Dally, N., Boughton, B. A., Woo, Y. H., Gao, G., Schijlen, E. G. W. M., Guo, X., Momin, A. A., Negrão, S., Al-Babili, S.,Gehring, C., Roessner, U., Jung, C., Murphy, K., Arold, S. T., Gojobori, T., Linden, C. G. V. D., Van Loo, E. N., Jellen, E. N., Maughan, P. J. & Tester, M. 2017. The genome of Chenopodium quinoa. Nature, 542, 307–312.

Kajitani, R., Toshimoto, K., Noguchi, H., Toyoda, A., Ogura, Y., Okuno, M., Yabana, M., Harada, M., Nagayasu, E., Maruyama, H., Kohara, Y., Fujiyama, A., Hayashi, T. & Itoh, T. 2014. Efficient de novo assembly of highly heterozygous genomes from whole-genome shotgun short reads. Genome Research, 24, 1384–1395.

Kingan, S. 2016. HomolContigsByAnnotation [Online]. Available: https://github.com/skingan/HomolContigsByAnnotation [Accessed 2017].

Koboldt, D. C., Zhang, Q., Larson, D. E., Shen, D., Mclellan, M. D., Lin, L., Miller, C. A., Mardis, E. R.,Ding, L. & Wilson, R. K. 2012. VarScan 2: somatic mutation and copy number alteration discovery in cancer by exome sequencing. Genome Res, 22, 568–76.

Koren, S., Walenz, B. P., Berlin, K., Miller, J. R., Bergman, N. H. & Phillippy, A. M. 2017. Canu: scalable and accurate long-read assembly via adaptive k-mer weighting and repeat separation. Genome Research, 27, 722–736.

Korlach, J., Gedman, G., King, S., Chin, J., Howard, J., Cantin, L. & Jarvis, E. D. 2017. De Novo PacBio long-read and phased avian genome assemblies correct and add to genes important in neuroscience research. bioRxiv.

Krzywinski, M. I., Schein, J. E., Birol, I., Connors, J., Gascoyne, R., Horsman, D., Jones, S. J. & Marra, M. A. 2009. Circos: An information aesthetic for comparative genomics. Genome Research.

Kurtz, S., Phillippy, A., Delcher, A. L., Smoot, M., Shumway, M., Antonescu, C. & Salzberg, S. L. 2004. Versatile and open software for comparing large genomes. Genome Biology, 5, R12–R12.

Lamesch, P., Berardini, T. Z., Li, D., Swarbreck, D., Wilks, C., Sasidharan, R., Muller, R., Dreher, K., Alexander, D. L., Garcia-Hernandez, M., Karthikeyan, A. S., Lee, C. H., Nelson, W. D., Ploetz, L.,Singh, S., Wensel, A. & Huala, E. 2012. The Arabidopsis Information Resource (TAIR): improved gene annotation and new tools. Nucleic Acids Research, 40, D1202–D1210.

Li, H. 2013. Aligning sequence reads, clone sequences and assembly contigs with BWA-MEM. ARXIV.

Loman, N. J., Quick, J. & Simpson, J. T. 2015. A complete bacterial genome assembled de novo using only nanopore sequencing data. Nat Meth, 12, 733–735.

Olson, N. D., Lund, S. P., Colman, R. E., Foster, J. T., Sahl, J. W., Schupp, J. M., Keim, P., Morrow, J. B., Salit, M. L. & Zook, J. M. 2015. Best practices for evaluating single nucleotide variant calling methods for microbial genomics. Frontiers in Genetics, 6, 235.

Pryszcz, L. P. & Gabaldon, T. 2016. Redundans: an assembly pipeline for highly heterozygous genomes. Nucleic Acids Res, 44, e113.

Pryszcz, L. P., NéMeth, T., Gácser, A. & GabaldóN, T. 2014. Genome Comparison of Candida orthopsilosis Clinical Strains Reveals the Existence of Hybrids between Two Distinct Subspecies. Genome Biology and Evolution, 6, 1069–1078.

Quinlan, A. R. & Hall, I. M. 2010. BEDTools: a flexible suite of utilities for comparing genomic features.Bioinformatics, 26, 841–842.

Safonova, Y., Bankevich, A. & Pevzner, P. A. 2015. dipSPAdes: Assembler for Highly Polymorphic Diploid Genomes. J Comput Biol, 22, 528–45.

Schwessinger, B., Sperschneider, J., Cuddy, W. S., Garnica, D. P., Miller, M. E., Taylor, J. M., Dodds, P. N., Figueroa, M., Park, R. F. & Rathjen, J. P. 2018. A Near-Complete Haplotype-Phased Genome of the Dikaryotic Wheat Stripe Rust Fungus Puccinia striiformis f. sp. tritici Reveals High Interhaplotype Diversity. mBio, 9.

SimãO, F. A., Waterhouse, R. M., Ioannidis, P., Kriventseva, E. V. & Zdobnov, E. M. 2015. BUSCO:assessing genome assembly and annotation completeness with single-copy orthologs. Bioinformatics, 31, 3210–3212.

Small, K. S., Brudno, M., Hill, M. M. & Sidow, A. 2007. A haplome alignment and reference sequence of the highly polymorphic Ciona savignyi genome. Genome Biology, 8, R41–R41.

Vanburen, R., Wai, C. M., Ou, S., Pardo, J., Bryant, D., Jiang, N., Mockler, T. C., Edger, P. & Michael, T.P. 2018. Extreme haplotype variation in the desiccation-tolerant clubmoss Selaginella lepidophylla. Nat Commun, 9, 13.

Vinson, J. P., Jaffe, D. B., O’Neill, K., Karlsson, E. K., Stange-Thomann, N., Anderson, S., Mesirov, J. P., Satoh, N., Satou, Y., Nusbaum, C., Birren, B., Galagan, J. E. & Lander, E. S. 2005. Assembly of polymorphic genomes: algorithms and application to Ciona savignyi. Genome Res, 15, 1127–35.

Weisenfeld, N. I., Kumar, V., Shah, P., Church, D. M. & Jaffe, D. B. 2017. Direct determination of diploid genome sequences. Genome Research, 27, 757–767.

