## Supplementary Materials for "Purge Haplotigs: Synteny Reduction for Third-gen Diploid Genome Assemblies"

### Workflows for Purge Haplotigs and Analysis

#### Purge Haplotigs Validation

The validation run using a simulated genome and simulated PacBio subreads is archived here: https://doi.org/10.5281/zenodo.1042847. The files and commands used to simulate the reads, and generate and curate the assembly are included.

#### Purge Haplotigs Case Study: Running Purge Haplotigs

- *Arabidopsis thaliana*

```
$ purge_haplotigs readhist cvi-0_col-0_f1.bam

$ purge_haplotigs contigcov -i cvi-0_col-0_f1.bam.gencov -o
  coverage_stats.csv -l 10 -m 90 -h 170

$ purge_haplotigs purge -g p_h_ctg.fasta -c coverage_stats.csv -b
  cvi-0_col-0_f1.bam -t 16 -a 60
```

- *Clavicorona pyxidata*

```
$ purge_haplotigs readhist cpyxidata.alignmentset.bam

$ purge_haplotigs contigcov -i cpyxidata.alignmentset.bam.gencov -o
  coverage_stats.csv -l 10 -m 60 -h 150

$ purge_haplotigs purge -g p_h_ctgs.fasta -c coverage_stats.csv -b
  cpyxidata.alignmentset.bam -t 12 -a 60
```

- *Vitis vinifera L. Cv.* Cabernet Sauvignon

```
$ purge_haplotigs readhist Vvinifera_cabsauv.bam

$ purge_haplotigs contigcov -i Vvinifera_cabsauv.bam.gencov -o
  coverage_stats.csv -l 10 -m 80 -h 170

$ purge_haplotigs purge -g p_h_ctg.fasta -c coverage_stats.csv -b
  Vvinifera_cabsauv.bam -t 16 -a 60 -wind_len 10000 -wind_step 5000
```

- *Taeniopygia guttata*

```
$ purge_haplotigs readhist aligned.bam

$ purge_haplotigs contigcov -i aligned.bam.gencov -o
  coverage_stats.csv -l 10 -m 65 -h 150

$ purge_haplotigs purge -g p_a_ctgs.fasta -c coverage_stats.csv -b
  aligned.bam -t 16 -a 60
```

Purge Haplotigs Case Study: Illumina Paired End Short Read Mapping

- Example workflow (same protocol used for all)
  - Create BWA index

```
$ bwa index genome.fasta
```

  - Map the reads

```
$ bwa mem genome.fasta -t 16 '< zcat reads_R1.fastq.gz' '< zcat
  reads_R1.fastq.gz' | samtools view -bS -q 10 - > aligned.bam
```

  - Sort

```
$ samtools sort -@ 8 -m 1G -o aligned.s.bam aligned.bam
```

  - Deduplicate

```
$ samtools rmdup aligned.s.bam aligned.sd.bam
```

  - Remove clipped and discordant reads

```
$ samtools view -h -f 0x2 aligned.sd.bam | awk '$6 !~ /H|S/{print}'
  | samtools view -bS - > aligned.sdc.bam
```

  - Get simple stats on mapping

```
$ samtools flagstat aligned.sdc.bam
```

  - Call variants

```
$ samtools mpileup -f genome.fasta aligned.sdc.bam | java -jar
  Varscan.jar mpileup2snp --p-value 0.001 | gzip - > aligned.SNPs.tsv.gz
```

  - Filter variants (heterozygous only, read-depth between 20 and 200 for *C. pyxidata* or 10 and 100 for *A. thaliana* and *T. guttata*)

```
$ zcat aligned.SNPs.tsv.gz | perl -e 'while(<>){my@l=split(/\s+/,$_);my
  @d=split(/:/,$l[4]);($l[7]==1)&&($d[1]>20)&&($d[1]<200)&&(print $_);}'
  | gzip - > aligned.SNPs.filt.tsv.gz
```

#### Purge Haplotigs Case Study: Circos Plots

The workflow example and control file examples are available at:
https://bitbucket.org/mroachawri/read_snp_circos_eg

#### Purge Haplotigs Case Study: BUSCO Analysis

- *A. thaliana*

```
$ python run_BUSCO.py -i cns_h_ctg.fasta -o cns-h-ctg -l
  /datasets/embryophyta_odb9/ -m genome -c 16 -sp arabidopsis
```

- *C. pyxidata*

```
$ python run_BUSCO.py -i cns_h_ctg.fasta -o cns-h-ctg -l
  /datasets/basidiomycota_odb9/ -m genome -c 24 -sp coprinus
```

- *V. vinifera L. Cv.* Cabernet Sauvignon

```
$ python run_BUSCO.py -i cns_h_ctg.fasta -o cns-h-ctg -l
  /datasets/embryophyta_odb9/ -m genome -c 24 -sp arabidopsis
```

- *T. guttata*

```
$ python run_BUSCO.py -i h_ctgs.fasta -o cns-h-ctg -l
  /datasets/aves_odb9/ -m genome -c 24 -sp human
```

#### Purge Haplotigs Case Study: MUMmer Alignments and Dotplots

- Example workflow
  - Calculate alignments between reference (`ref.fa`) and query (`query.fa`)

```
$ nucmer -t 16 ref.fa query.fa -p ref-query
```

  - Filter alignments

```
$ delta-filter -1 ref-query.delta > ref-query.1.delta
```

  - To make a dotplot (the mummerplot script was modified to reduce the size of the points and give thinner lines when plotting)

```
$ mumemrplot --fat --png --large ref-query.1.delta -p ref-query
```

  - To get the bases in ref covered by alignments from query for calculating coverage

```
$ show-coords -H ref-query.1.delta | awk '{ len += $7 }END{ print len }'
```
