## Supplementary Materials for "Purge Haplotigs: Synteny Reduction for Third-gen Diploid Genome Assemblies"

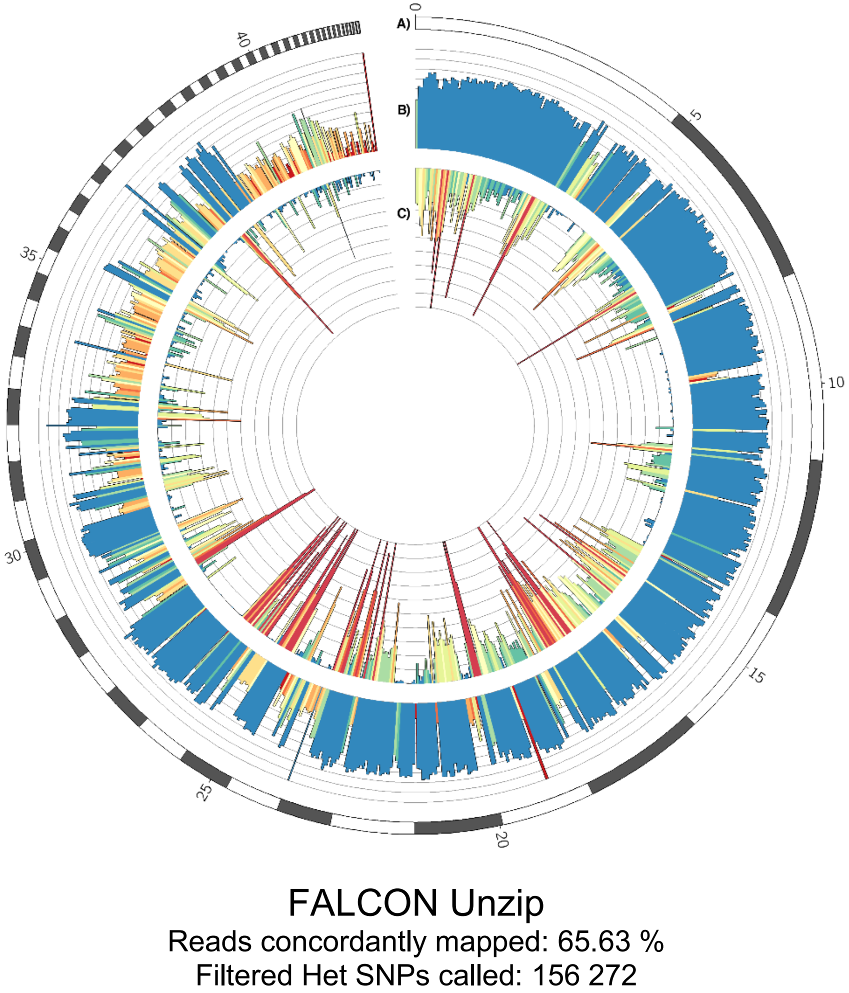


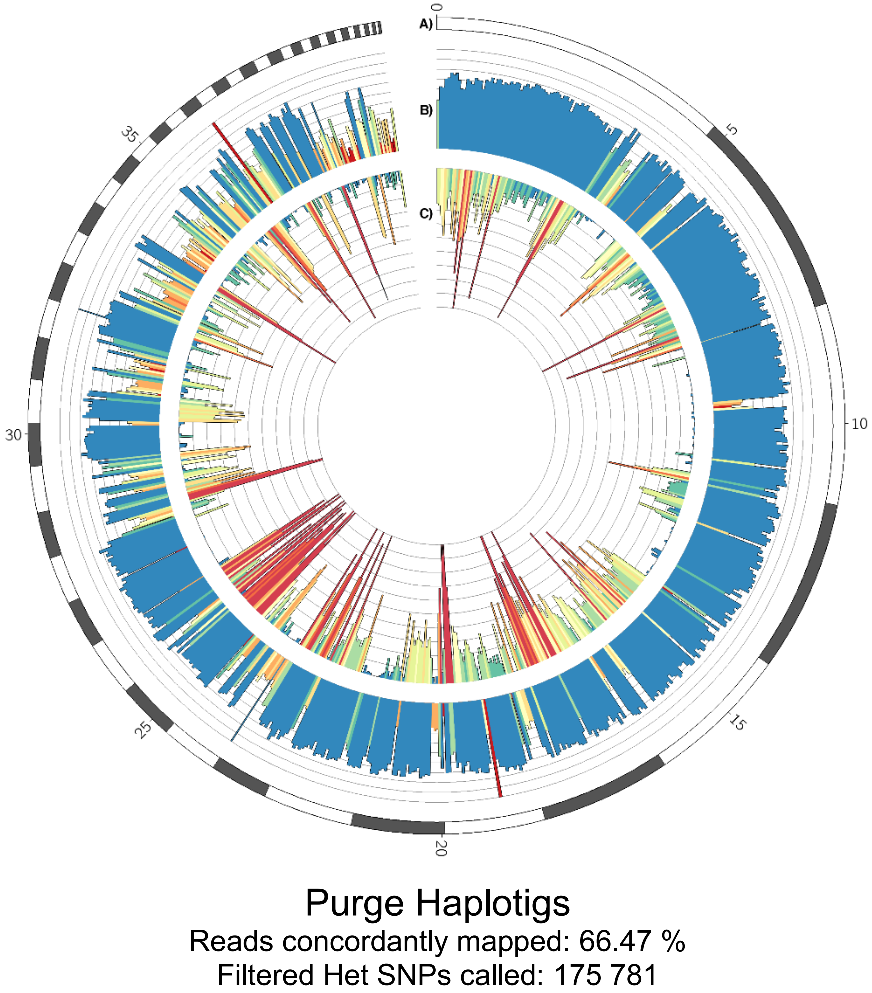


**Circos plots for *Clavicorona pyxidata* haploid assemblies.** Short paired end reads were mapped (SRA accession: SRR1800147) and SNPs were called for the draft FALCON Unzip assembly (LEFT) and the assembly curated with Purge Haplotigs (RIGHT). The tracks shown in the circos plots are: **A)** Contigs (ordered by length), **B)** Read-depth histogram (reads per genome window), **C)** SNP density (SNPs per genome window).

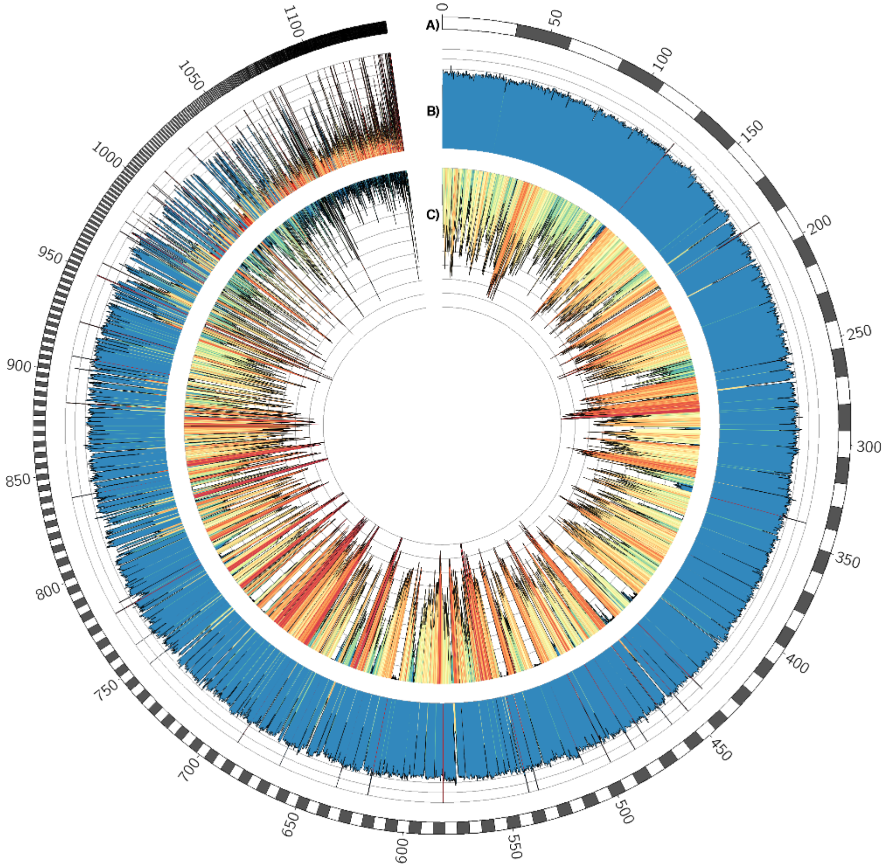


### FALCON Unzip

Reads concordantly mapped: 86.08 %
Filtered Het SNPs called: 8 931 286

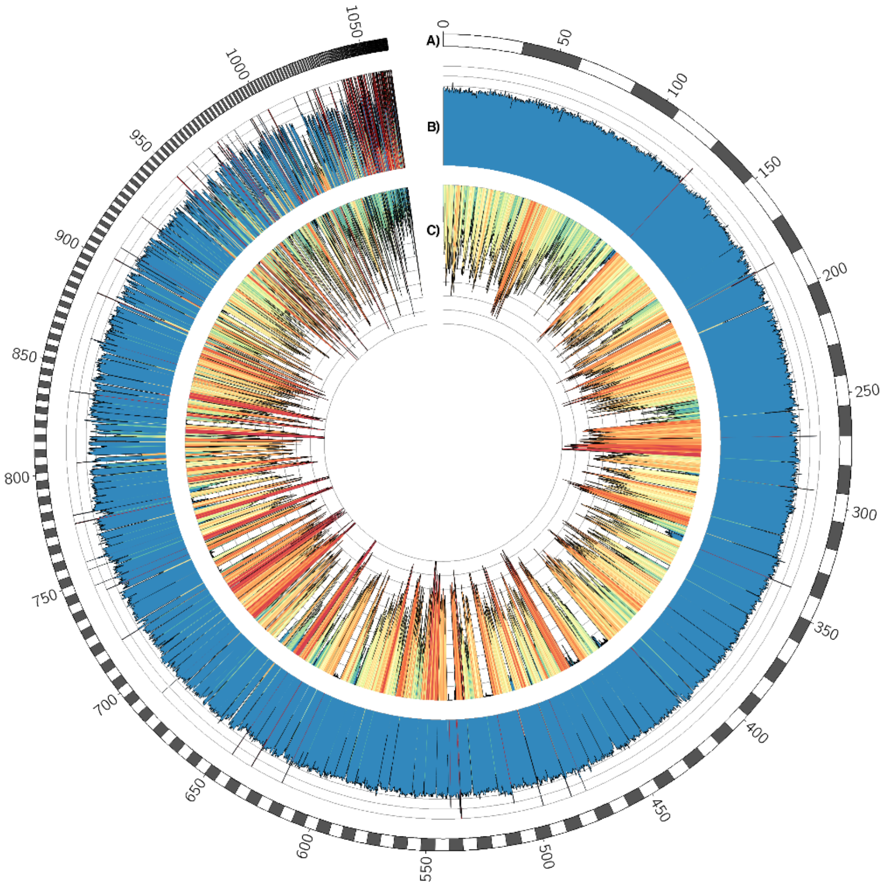


### Purge Haplotigs

Reads concordantly mapped: 86.68 %
Filtered Het SNPs called: 9 125 759

**Circos plots for *Taeniopygia guttata* haploid assemblies.** Short paired end reads were mapped (SRA accession: ERR1013157) and SNPs were called for the draft FALCON Unzip assembly (LEFT) and the assembly curated with Purge Haplotigs (RIGHT). The tracks shown in the circos plots are: **A)** Contigs (ordered by length), **B)** Read-depth histogram (reads per genome window), **C)** SNP density (SNPs per genome window).
