## Supplementary Materials for "Purge Haplotigs: Synteny Reduction for Third-gen Diploid Genome Assemblies"

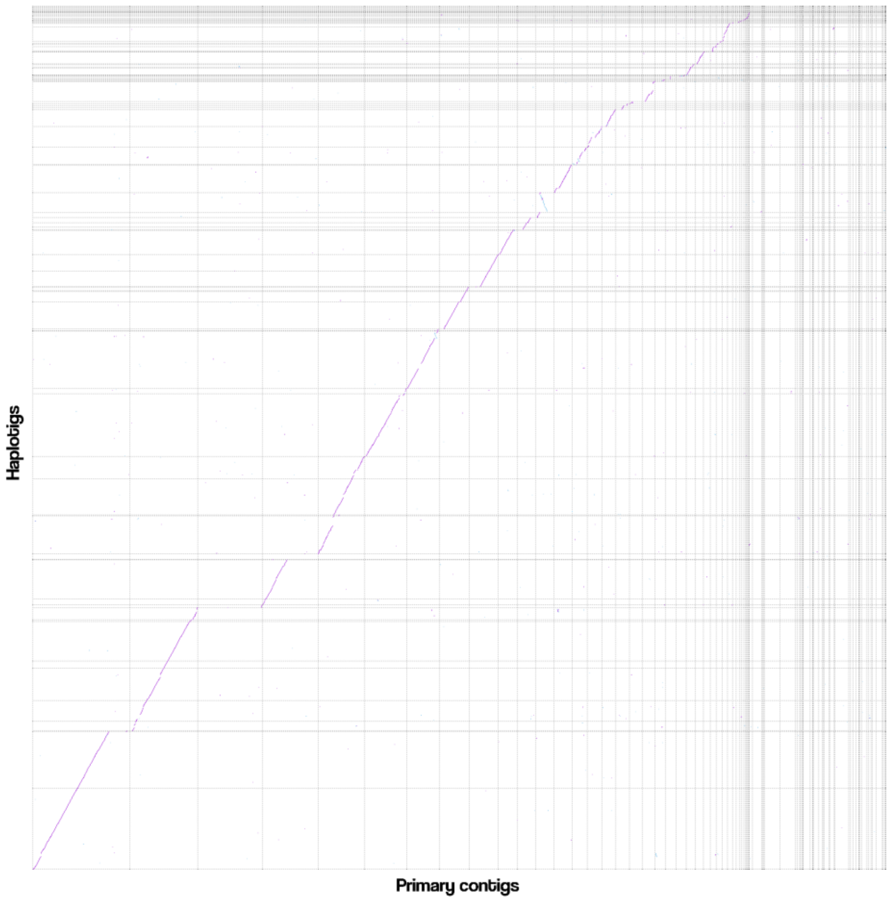


FALCON Unzip
Primary contig coverage: 49.7 %

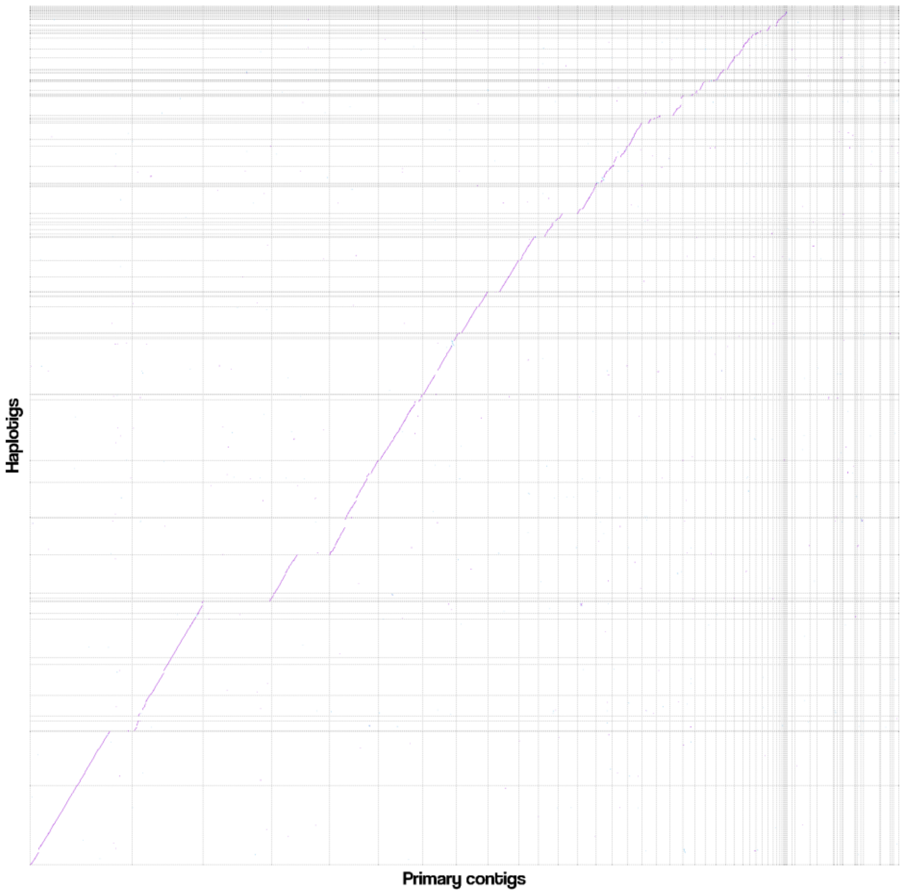


Purge Haplotigs
Primary contig coverage: 52.9 %

**Dotplots for *Clavicorona pyxidata* assemblies.** Haplotigs were aligned to primary contigs, filtered for one-to-one best alignments, coverage of the primary contigs by haplotigs calculated, and dotplots were laid out by longest alignments.

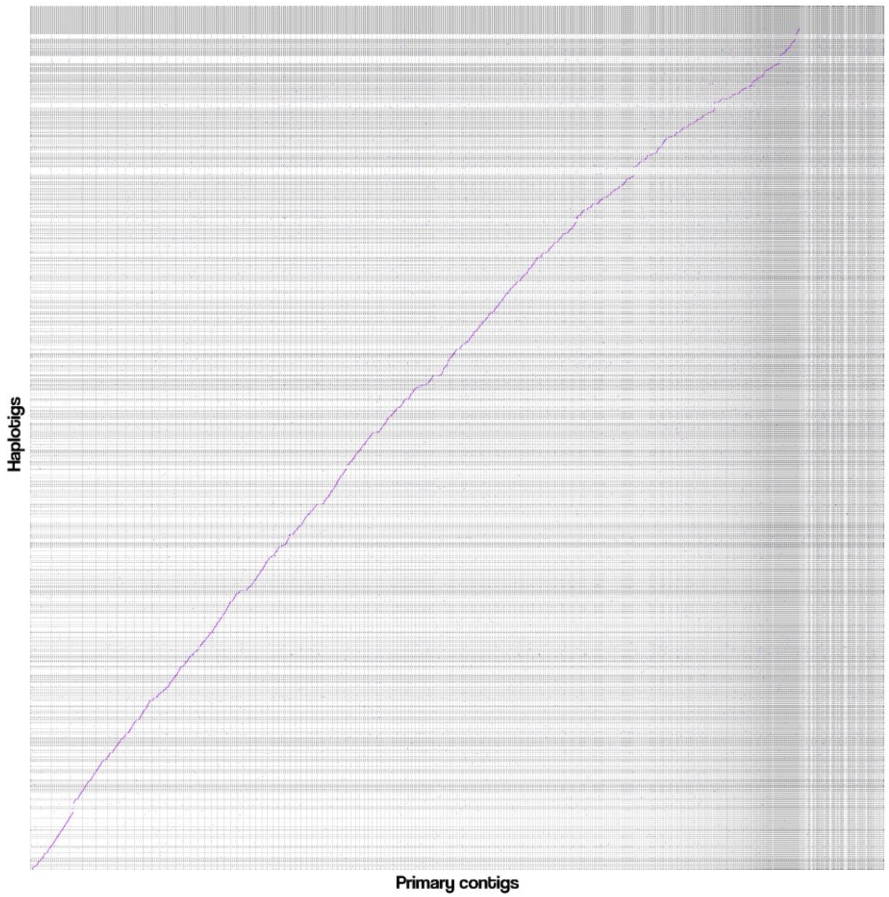


FALCON Unzip
Primary contig coverage: 48.5 %

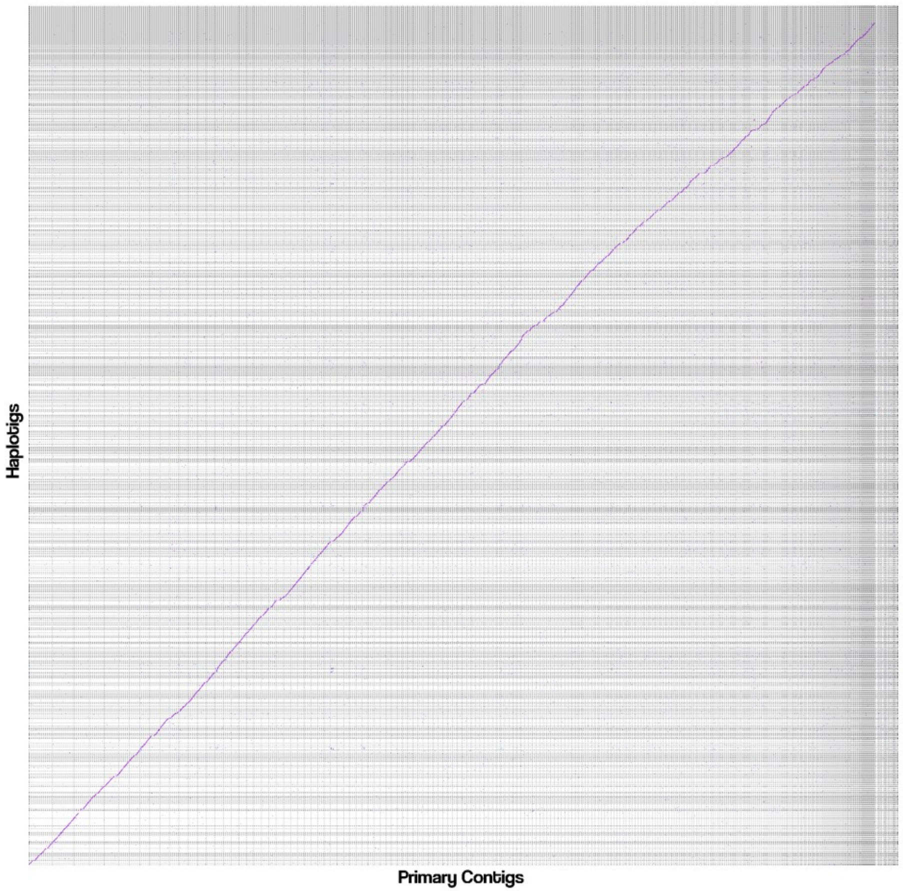


Purge Haplotigs
Primary contig coverage: 64.3 %

**Dotplots for *Vitis vinifera L. Cv.* Cabernet Sauvignon assemblies.** Haplotigs were aligned to primary contigs, filtered for one-to-one best alignments, coverage of the primary contigs by haplotigs calculated, and dotplots were laid out by longest alignments.

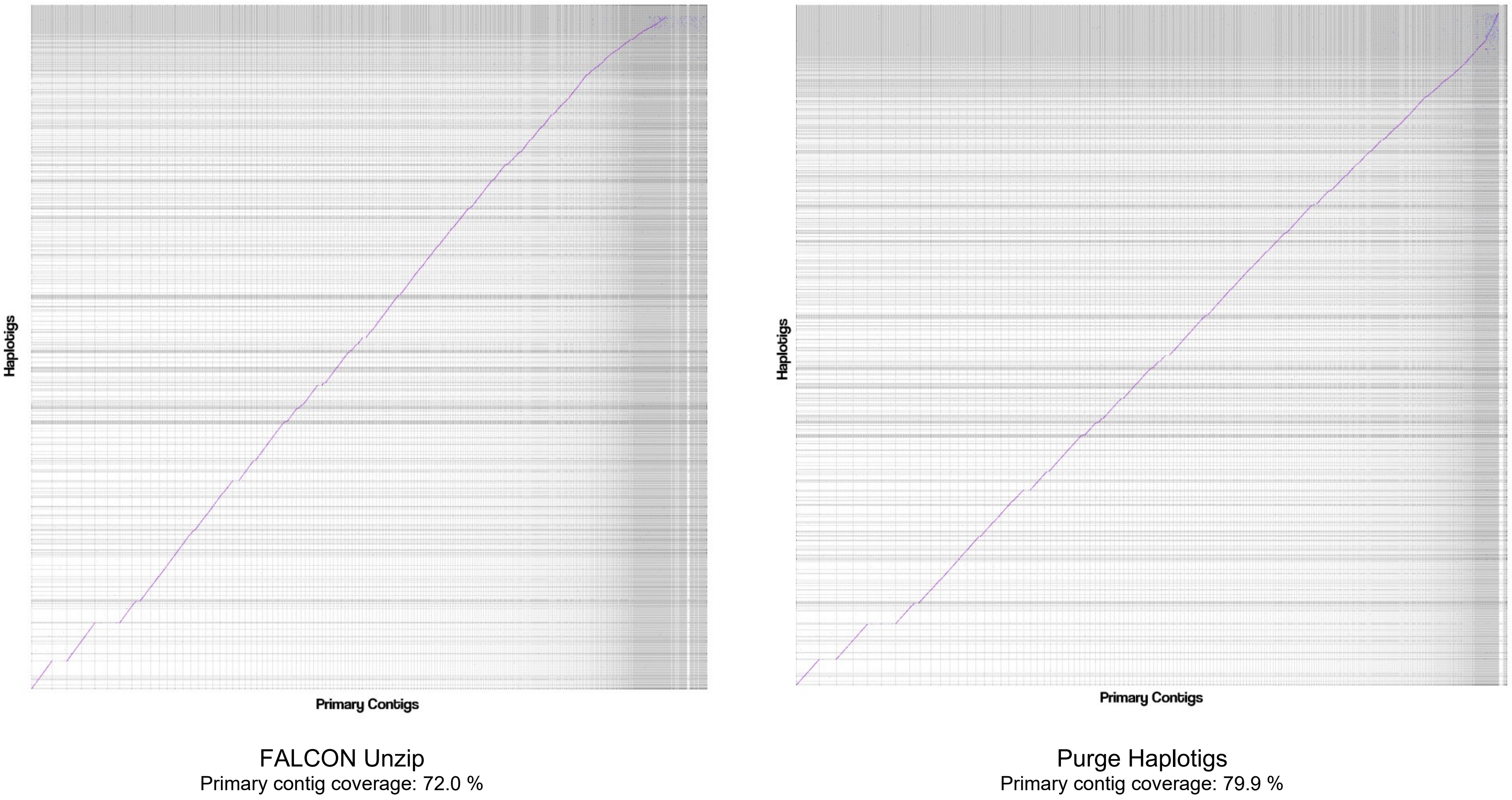


**Dotplots for *Taeniopygia guttata* assemblies.** Haplotigs were aligned to primary contigs, filtered for one-to-one best alignments, coverage of the primary contigs by haplotigs calculated, and dotplots were laid out by longest alignments.
